# Using developmental dynamics for evolutionary prediction and control

**DOI:** 10.1101/2023.11.03.565446

**Authors:** Lisandro Milocco, Tobias Uller

## Abstract

Understanding, predicting, and controlling the phenotypic consequences of genetic and environmental change is essential to many areas of fundamental and applied biology. In evolutionary biology, the generative process of development is a major source of organismal evolvability that constrains or facilitates adaptive change by shaping the distribution of phenotypic variation that selection can act upon. While the complex interactions between genetic and environmental factors during development may appear to make it impossible to infer the consequences of perturbations, the persistent observation that many perturbations result in similar phenotypes indicates that there is a logic to what variation is generated. Here, we show that a general representation of development as a dynamical system can reveal this logic. We build a framework that allows to predict the phenotypic effects of perturbations, and conditions for when the effects of perturbations of different origin are concordant. We find that this concordance is explained by two generic features of development, namely the dynamical dependence of the phenotype on itself and the fact that all perturbations must be funneled by the same developmental process. We apply our theoretical results to classical models of development and show that it can be used to predict the evolutionary response to selection using information of plasticity, and to accelerate evolution in a desired direction. The framework we introduce provides a way to quantitatively interchange perturbations, opening a new avenue of perturbation design to control the generation of variation, and thus evolution.

## Introduction

A complete theory of organismal evolution requires a theory of phenotypic variation, a theory of natural selection, and a theory of heredity. While tremendous advances have been made in the last century to understand the two latter pillars of Darwinian evolution, a theory for the generation of phenotypic variation remains elusive.

The process that generates variation in morphology, physiology, and behavior is known as development in the broad sense (Gilbert and Barresi 2016). Notoriously complex and non-linear interactions between genes, cells, tissues and environmental factors during development make it difficult to grasp the phenotypic consequences of genetic and environmental perturbations. Indeed, the diversity and complexity of developmental systems could be taken as evidence that *a priori* inference of the consequences of perturbations rarely will be feasible. A pessimistic conclusion is therefore that the best one could hope for is to demonstrate that generative processes in principle can impact evolutionary trajectories (Rice 2002, Morrissey 2015, Gonzalez-Forero 2023), while studies that demonstrate *how* development affects evolution will remain a collection of idiosyncratic case studies (Beldade et al. 2002, Brakefield 2006, Galis et. al 2010). This perception that generative processes are intrinsically unpredictable, and that selection is the only reliable force in evolution is also reflected in biotechnology and medicine, where attempts to direct evolutionary processes emphasize control over selective regimes rather than control over generative processes.

In this paper, we provide a more optimistic perspective by addressing a particular problem concerning the generation of variation, and its implications for evolution: the relationship between the phenotypic effects of genetic and environmental perturbation. Genetic and environmental effects on phenotypic variation have often been considered independent, as implicitly assumed when environmental effects are modelled as the uncorrelated residuals of a linear regression of phenotype on genotype in quantitative genetics (Lynch and Walsh 1998). However, since both genetic and environmental perturbations are channeled through the same developmental system, it is unlikely that this assumption generally holds true (Cheverud 1988, West-Eberhard 2003). It is indeed well known that environmental change occasionally induces phenotypes that resemble genetic mutants (e.g., melanism in butterflies, Nijhout 1984) and it has been shown that plastic responses are biased towards phenotype dimensions with high additive genetic variation (Noble et al. 2019), but the existing body of work is mostly a collection of empirical observations.

If genes and environments are equivalent, or interchangeable, as sources of phenotypic variation, this could have important consequences for understanding and predicting evolution, and eventually controling it. In particular, gene-environment interchangeability implies that there is a formal connection between evolvability and plasticity. Evolvability can be defined as the capacity to generate phenotypic variation in response to genotypic variation (Kirschner and Gerhart 1998), while plasticity refers to the same capacity for phenotypic variation in response to environmental variation.

If genetic and environmental perturbations are interchangeable, the evolution of plasticity may shape evolvability and *vice versa*, and information of one can reveal features of the other (Chevin et al. 2022). Previous theoretical work has suggested that such a relationship between plasticity and evolvability does exist (Wagner and Altenberg 1996, Ancel and Fontana 2000, Espinosa-Soto et al. 2011, Draghi and Whitlock 2012, Furusawa and Kaneko 2015, van Gestel and Weissing 2016, Brun-Usan et al. 2021), but there is no general framework to explicitly define the conditions for when this relationship should be expected, or to study its evolutionary implications. Such understanding would enable the design combinations of perturbations to drive the developmental system to a desired state, thus controlling the generation of variation.

The aim of this paper is to introduce a conceptual framework to understand when genetic and environmental perturbation will cause shifts in phenotype in similar directions in trait space. We illustrate this phenomenon of alignment using *in silico* experiments of reaction diffusion models and gene regulatory networks. We show how the theory can be used (i) to predict the concordance of phenotypic effects of perturbations of different origins, (ii) to estimate the effects of mutations on the phenotype, (iii) to infer evolvability using information of plasticity, and (iv) to accelerate evolution in a desired direction. This ability to convert information from plastic responses into information about evolutionary potential, and vice versa, could have applications in diverse areas concerned with the phenotype, including developing solutions to environmental and societal challenges using biotechnological engineering.

## Results

The results are presented in sections. First, we introduce a general representation of development as a dynamical system. Second, we develop the formalism to study the phenotypic effect of a single perturbation. Third, we study the alignment between perturbations of different origins (e.g., genetic and environmental). Finally, we apply the theoretical framework to classical models of development, namely reaction-diffusion models and gene regulatory networks, and show how it can be used for evolutionary understanding, prediction and even control.

### A general representation of development as a dynamical system

Mathematical models of development usually consist of a representation of the phenotype and a set of rules of how this phenotype changes through developmental time, for example, through the interaction among different components of the system. Examples of such models include reaction-diffusion models (e.g., Kondo and Miura 2010), gene regulatory networks (e.g., Wagner 1994), and models of morphogenesis (e.g., Salazar-Ciudad and Jernvall 2010). These models are commonly given mathematically as differential equations which are numerically integrated over time to simulate a developmental trajectory, which is the change in the phenotypic values through developmental time. Following this body of work (Lewontin 1983, Alberch 1991), we take the general representation of development given by:

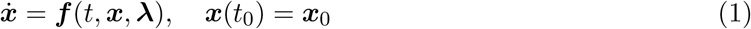

where ***x*** = (*x*_1_, *x*_2_, …, *x*_*n*_) is a vector composed of *n* variables that we refer to as *states*, with each state *x*_*i*_ representing a different aspect of the phenotypes that is relevant to describe the systems behavior through developmental time (e.g., the expression level of a given gene); 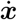 is the time derivative of ***x***, which gives the temporal change in the states; *t* is developmental time; ***f*** is a developmental function determining the rules of how the states change in time; ***x***_0_ are the state values at initial time *t*_0_, known as the initial conditions; and ***λ*** = (*λ*_1_, *λ*_2_, …, *λ*_*p*_) are developmental parameters, which can be genetic or environmental (e.g., the affinity of a cofactor modulating downstream gene expression, or temperature during developmental time).

Equation (1) captures two central properties of development which will be important to derive the results presented later. The first of these central aspects is that development depends at each step on the preexisting phenotype. This is mathematically captured by the fact that the change in the states at each time, given by 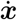, is itself a function of the states ***x*** at that time. This means that the phenotype at any given time is both the effect of earlier and the cause of later developmental changes. This feedback of the phenotype on itself makes development a *dynamical* phenomenon rather than a *static* one (or *historical* rather than *programmatic*, Stent 1985, West-Eberhard 2003), where the ways in which the phenotype can and cannot change at a given time depend on the state of the phenotype at that time. Examples of this historicity of development include the sequential determination of cell fate (Bassett and Wallace 2012) and sensitivity windows, where the same perturbation results in a phenotypic effect only for responsive phenotypes at specific times during development (Burggren and Mueller 2015).

The second important aspect of development highlighted by equation (1) is that changes in any of the developmental parameters ***λ*** have an effect on the states ***x*** through the same function ***f***. In other words, any perturbation in the developmental parameters has to be channeled through the same developmental pathways to result in a change in the states. As we show below, this *funneling* (Cheverud 1988, West-Eberhard 2003) is fundamental for the alignment of the effects of perturbations with different origins.

### The effect of a perturbation on one developmental parameter

We are interested in studying how a given developmental trajectory is affected by a perturbation in one of the developmental parameters. We begin with a system with a single developmental parameter (i.e., ***λ*** = *λ*), and we extend the results to multiple parameters later. Further, we will assume that the developmental function ***f*** is smooth, having continuous fist partial derivatives.

The study of perturbations is always comparative: perturbations must be studied with respect to an unperturbed reference. We thus need to define a reference developmental trajectory from which to study deviations from. We define *λ*^*^ as the reference value for the developmental parameter (i.e., corresponding to an organism with the wild-type genotype in standard environmental conditions). The developmental trajectory of the reference, unpertubed developmental system is thus given as the unique solution of equation (1) for *λ* = *λ*^*^, which we denote ***x***(*t, λ*^*^), as shown in Figure 1.

**Figure 1:**
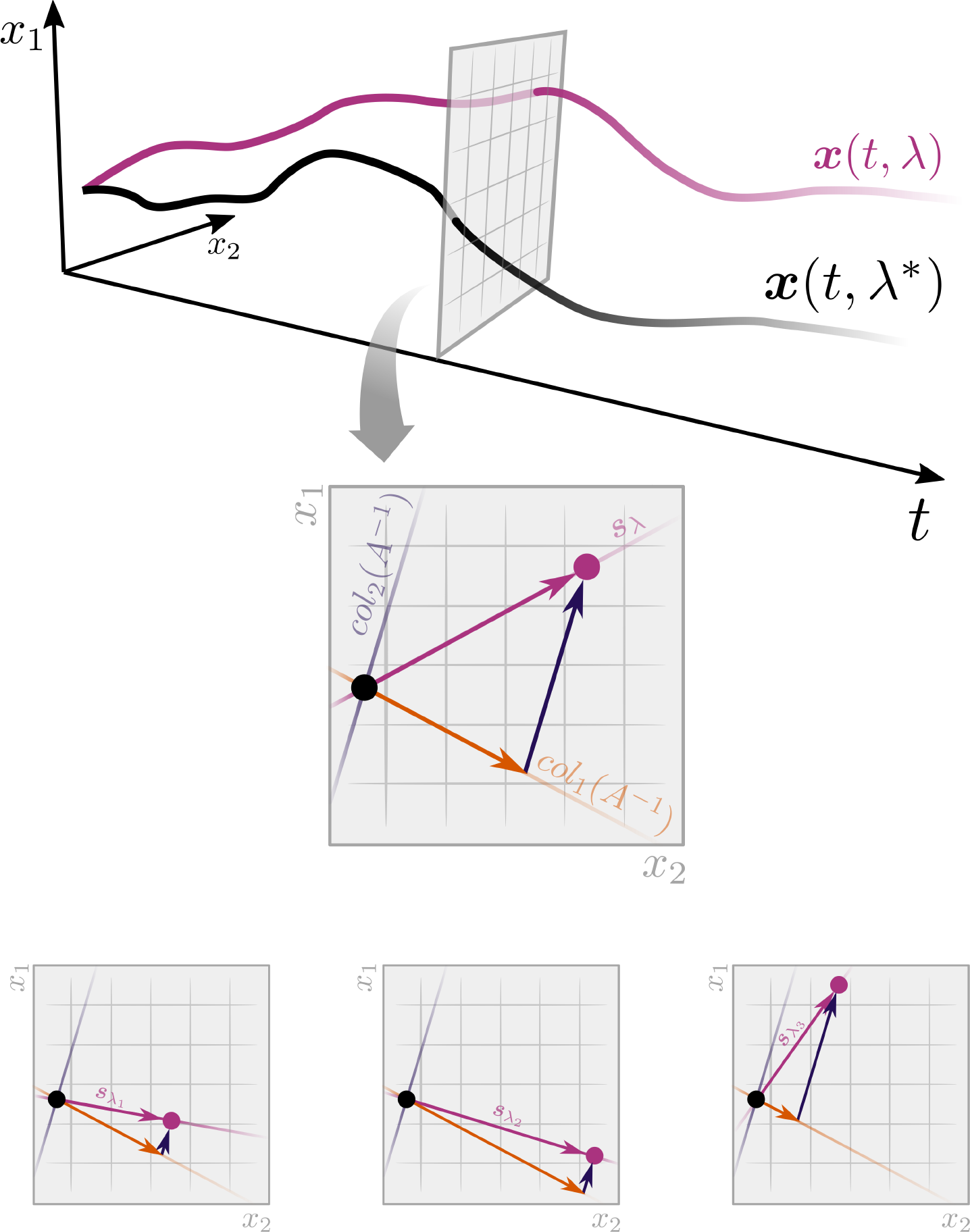
A general framework to study the phenotypic effects of perturbations of different origins. On top, the reference developmental trajectory ***x***(*t, λ*^*\**^) in black and the perturbed trajectory ***x***(*t, λ*) in purple through developmental time *t*. The panel in the middle shows that, at any given time, the effect of the perturbation on the trajectory, given by the sensitivity vector ***s***_*λ*_(*t*) is a linear combination of the columns of the matrix *A*^*−*1^(*t, λ*^*\**^). The three panels at the bottom show the sensitivity vectors for different perturbations at a given developmental time are linear combinations of the columns of the same 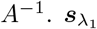 and 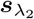 are largely aligned, but not 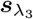.

We are interested in the direction in which the reference developmental trajectory will change when we introduce a small perturbation to *λ*^*^. This type of study is knows as *sensitivity analysis* in dynamical systems theory (Khalil 2002). The direction of change is given by

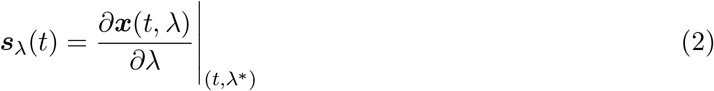

The vector ***s***_*λ*_(*t*) is known as the *sensitivity vector* (or *function*, Khalil 2002), and it is a vector of length *n* containing the partial derivatives of the states *x*_1_, *x*_2_, …, *x*_*n*_ with respect to the parameter *λ*, evaluated at the reference value *λ*^*^ and at time *t*. This vector then tells us how we expect the states to change for a small change in the developmental parameter at each time *t*. For small perturbations, we can predict the perturbed developmental trajectory using:

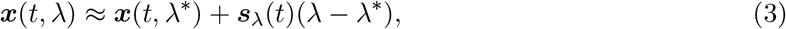

which tells us that the perturbed developmental trajectory ***x***(*t, λ*) will differ from the reference, unperturbed trajectory ***x***(*t, λ*^*^) by an amount proportional to the difference *λ* − *λ*_0_, with direction determined by the vector ***s***_*λ*_(*t*). Equation (3) resembles a first order Taylor approximation, and is only locally valid (i.e., for values of *λ* close to *λ*^*^).

Calculating ***s***_*λ*_(*t*) is not straight-forward since we do not know the explicit relationship between the states ***x*** and the parameter *λ*. We show in *Appendix A* (see also Khalil 2002) that ***s***_*λ*_(*t*) can be obtained as the the unique solution to

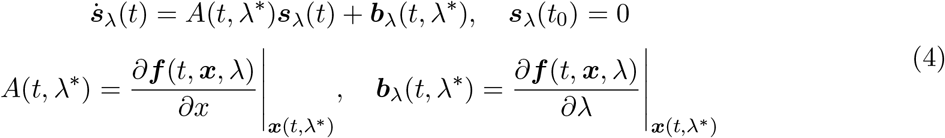

The matrix *A*(*t, λ*^*^) is known as the Jacobian matrix and summarizes the relationship between ***f*** and ***x***. Note that this Jacobian does not depend on what parameter *λ* is perturbed. The relationship between ***f*** and *λ* is captured by the vector ***b***_*λ*_(*t, λ*^*^). In this way, if we know the function ***f***, then we can calculate *A*(*t, λ*^*^) and ***b***_*λ*_(*t, λ*^*^), and jointly solve numerically equations (1) and (4) to obtain ***s***_*λ*_(*t*), which is the vector of interest.

Under the assumption that the Jacobian is invertible, we can get the simplified expression

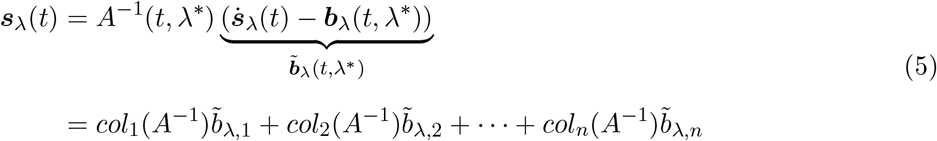

where *col*_*i*_(*A*^−1^) is the *i*-th column of *A*^−1^(*t, λ*^*^) and 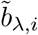 is the *i*-th element of vector 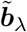. This means that the sensitivity vector at a given time ***s***_*λ*_(*t*) can be expressed as a linear combination of the columns of *A*^−1^(*t, λ*^*^) with weights determined by the elements of 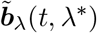. This result is shown graphically in Figure 1, and provides a basis to study alignment as explained in the next section. Note that 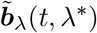 reduces to ***b***_*λ*_(*t, λ*^*^) if 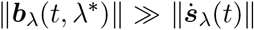, which is the case for an organisms that has reached steady state (e.g., adulthood).

### Alignment between the effects of perturbations of different origin

We now use the formalism introduced above to study the relationship between the phenotypic effects of perturbations of different origins. Given two developmental parameters, we say that their effects are totally aligned if the associated sensitivity vectors have an angle of 0°, meaning that the two perturbations result in phenotypic changes in exactly the same direction. More generally, we say that there is evidence of alignment if the two sensitivity vectors have an angle that is significantly smaller, at a given confidence level, than the distribution of angles between independent random vectors of the same dimension.

Figure 1 gives an example for three developmental parameters *λ*_1_, *λ*_2_ and *λ*_3_, which can correspond for example to the affinity of a cofactor modulating gene expression (genetic parameter), temperature and salinity (environmental parameters), respectively. The effects of modifying each of those parameters at time *t* is given by the vectors 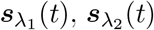 and 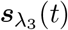, and the angles between them determine alignment. From equation (5), we know that all of these sensitivity vectors can be written as a linear combinations of the columns of the same matrix, the Jacobian *A*^−1^(*t*, ***λ***^*^), where each column is weighted by the elements of 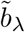 (note that we omit the arguments (*t, λ*^*^) when it is clear from context for readability). This provides sufficient conditions for alignment between the effects of different perturbations; if two perturbations have a dominant component in the direction of one of the columns of *A*^−1^(*t*, ***λ***^*^) (i.e., 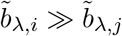for all *j* ≠ *i*), then these perturbations will be aligned.

The illustrative example in Figure 1 shows that 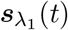 and 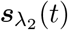 are largely aligned because they both have a large component in the direction of the first column and a small component in the direction of the second column of the Jacobian (i.e.,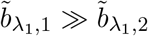 and 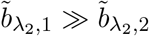). 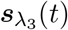 is not aligned with the other two vectors, since it has a large component in the second rather than first column (i.e., 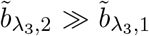). We note that this sufficient condition is not necessary for alignment. Indeed, there can be alignment according to our definition even if weights are not proportional when the Jacobian has columns that are similar to each other.

The two components of the sensitivity vectors, namely *A*^−1^(*t*, ***λ***^*^) and 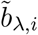, are related to the two key aspects of development highlighted by equation (1). The first of these aspects – the historicity of development – is related to matrix *A*^−1^(*t*, ***λ***^*^), the inverse of the Jacobian which summarizes the relationship between ***f*** and ***x***. This matrix determines the structure for phenotypic changes, since its columns provide the directions in which the phenotype is able to respond to a perturbation. These directions are independent of the nature of the perturbation itself and are determined by the capabilities of the responsive phenotype at that time. The second relevant aspect of development highlighted by equation (1) is the fact that all perturbations are funneled by the same developmental function ***f***. This is related to the other component of the sensitivity vector, the weights 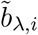. These weights summarize the relationship between ***f*** and ***λ***. If two developmental parameters affect the same aspects of ***f***, then there will be alignment. However, if two parameters affect distinct aspects of ***f*** (e.g., they affect two different developmental modules) then we should not expect alignment in general.

In the next sections, we apply this general framework to well-known models of development, and use it to make evolutionary prediction and control.

### Alignment in a reaction-diffusion model

Reaction–diffusion models are a set of models of pattern-formation that have been widely used to represent diverse developmental processes, including digit formation and hair follicle placement (Turing 1952, Sick et al. 2006, Kondo and Miura 2010, Green and Sharpe 2015). These models consist of a physical representation of the tissue and a set of molecules called morphogens. Morphogens diffuse and interact within the tissue, leading to the emergence of patterns as they accumulate in specific spatial regions. Here, we will use one of these reaction-diffusion models known as the Gray-Scott model (Gray and Scott 1990) to illustrate how we can study the alignment of phenotypic effects of different origin using the framework introduced in the previous sections.

The developmental function ***f*** for the Gray-Scott model is given by

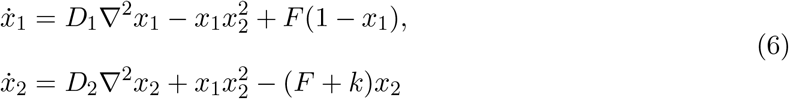

where *x*_1_ and *x*_2_ are the cellular concentrations of morphogen 1 and 2, respectively, *D*_1_ and *D*_2_ are their diffusion rates to neighboring cells, ∇^2^ is the Laplacian operator, *F* is the production rate of morphogen 1, and *k* is the rate of degradation of morphogen 2. We will study the alignment between the phenotypic effects of perturbing the developmental parameters *k* and *F*.

We represent a portion of embryonic tissue as a grid of 50 × 50 cells (details of the simulations are given in *Materials and Methods*). In each of these cells there is a given amount of the two morphogens, so the total number of states in the system is 50 × 50 × 2 = 5000. Diffusion of the morphogens occurs between neighboring cells. The simulation is done for a window of time of *t* = 5000 × *h* where *h* = 0.1 is the integration step. We start from initial conditions shown in Supplementary Figure 1, and use the reference parameters values 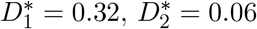, *k*^*^ = 0.06 and *F* ^*^ = 0.032. We calculate ***s***_*k*_(*t*) and ***s***_*F*_ (*t*) by jointly integrating equation (4).

Figure 2 shows that the angle between ***s***_*k*_(*t*) and ***s***_*F*_ (*t*) remains around 50°. This means that ***s***_*k*_(*t*) and ***s***_*F*_ (*t*) are partially aligned, since this angle is significantly smaller than the angle between random vectors of the dimension of the sensitivity vectors, which is 90°. This implies that the phenotypic effects of perturbing *k* and *F* should be similar.

**Figure 2:**
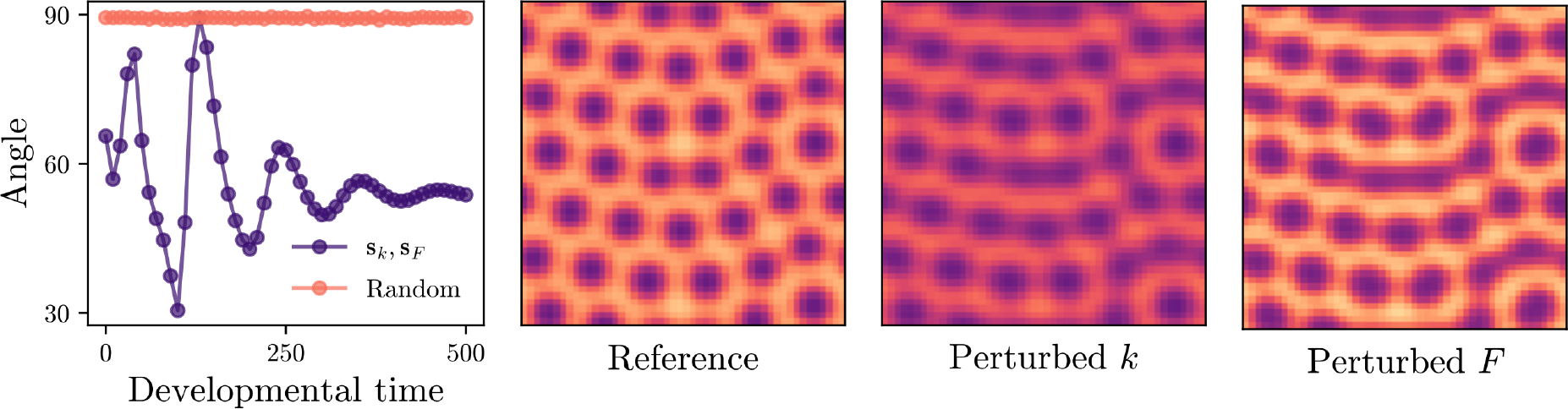
Alignment in a reaction-diffusion model. The panel on the left shows in purple the angle between ***s***_*F*_ (*t*) and ***s***_*k*_(*t*) through development time, and in orange the average angle between ***s***_*k*_(*t*) and 10 random vectors of the same dimension (one standard deviation is also plotted but covered by the dots). The angle between ***s***_*F*_ and ***s***_*k*_ is significantly smaller that the angle between random vectors, indicating alignment between the phenotypic effects of perturbing *k* and *F*. The three panels to the right show the phenotypes, plotted as the concentration of morphogen 1 with higher concentration in lighter color, for the reference developmental parameters, perturbed *k* and *F*, respectively, at developmental time 500. As indicated by the small angle between ***s***_*F*_ and ***s***_*k*_, the phenotypic effect of the perturbations is similar, resulting in *connected dots* as opposed to the *dotted* pattern in the reference.

We test the analytical prediction of alignment by simulating perturbed systems. We run simulations with small perturbations in the developmental parameters (i.e., *k* = *k*^*^ + Δ*k* and *F* = *F* ^*^ + Δ*F*), and compare the resulting phenotypes. Figure 2 shows that the phenotypic effects of either a decrease in *k* or an increase in *F* are largely aligned, resulting in the formation of *connected dots* rather than *dots* as in the reference, unperturbed phenotype. Note that since the angle between ***s***_*k*_(*t*) and ***s***_*F*_ (*t*) is not 0°, we should not expect the perturbed phenotypes to be identical.

### Alignment in a gene regulatory network

We now use the general framework to study the alignment between the phenotypic effects of perturbation with different origins in a gene regulatory network. In this section, we derive analytical results and test them using simulations. In the next section, we use the knowledge of alignment to connect plasticity and evolvability.

We use a common representation of gene regulatory networks found in the literature (Wagner 1994, Draghi and Whitlock 2012, Brun-Usan et al. 2021), where the states are given by ***x*** = (*x*_1_, *x*_2_, …, *x*_*n*_) representing the expression levels of *n* transcription factors that regulate each other’s expression. The function ***f*** giving the change in the states during development for this example is

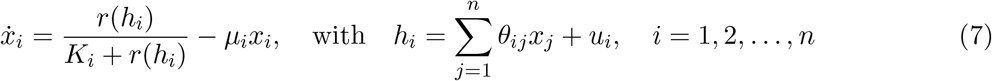

where *θ*_*ij*_ is the *ij*-th element of the matrix Θ, and gives the regulatory effect of gene *j* on the expression of gene *i* (i.e., *θ*_*ij*_ *>* 0, *θ*_*ij*_ *<* 0 and *θ*_*ij*_ = 0 represent activation, inhibition and no interaction, respectively). The expression of each gene can also be activated or inhibited by environmental inputs ***u*** = (*u*_1_, *u*_2_, …, *u*_*n*_). Gene expression follows Michaelis-Menten dynamics with coefficients *K*_*i*_, and gene product has a degradation rate given by *μ*_*i*_. In this way, the developmental parameters in this example are *θ*_*ij*_, *u*_*i*_, *K*_*i*_ and *μ*_*i*_ for all *i, j* = 1, 2, …, *n*. For the analyses in this section and the next using the gene-regulatory network, we study the steady state, which we consider maturity of the organism, where gene expression is no longer changing (i.e., 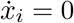 for all *i*). We use a bar to denote that a variable corresponds to the steady state (i.e., *Ā* is the Jacobian at the steady state).

In *Appendix B*, we obtain the Jacobian *Ā* and the weights 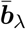, for each of the developmental parameters, and use them to calculate the sensitivity vector using equation (5). We find that for a given *i*, the sensitivity vectors 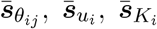, and 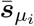 are always aligned (i.e., regardless of *j*) since the weight vectors 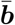 all have a non-zero value in the *i*-th position, and zeroes elsewhere. This means that, for example, a perturbation in the environmental input *u*_*i*_ will result in a phenotypic change that is in the same direction as a genetic change in any of the elements of the *i*-th row of Θ. In particular, this phenotypic effect will occur in the direction of vector *col*_*i*_*Ā*^−1^.

We test the analytical predictions by simulating gene networks of 5 genes and initial concentrations of 0.1 for all genes. We begin by using the sensitivity vectors to predict the phenotypic effects of mutations, which are changes in the elements of the interaction matrix Θ. For this, we generate 100 random gene regulatory networks each with a different reference interaction matrix 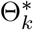. For each network *k*, we generate 20 mutants by modifying one element of 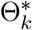. We predict the effect of these mutations using equation (3) as 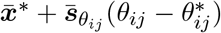, where 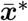 is the steady state of the unperturbed system and 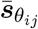 is as obtained in *Appendix B*. We then compare this prediction with the actual simulated steady state for the mutants.

Figure 3.a. shows that the formalism based on sensitivity vectors predicts the effect of mutations on the phenotype. As expected, the error in the prediction goes to zero as the perturbations become smaller. Perturbations smaller than 20% in the parameters have median relative error smaller than 3% in the prediction of their phenotypic effect. The predictions for this class of network is robust to larger perturbation, with perturbations in the range of 80-100% in the parameters still resulting in predictions with less than 30% median error.

**Figure 3:**
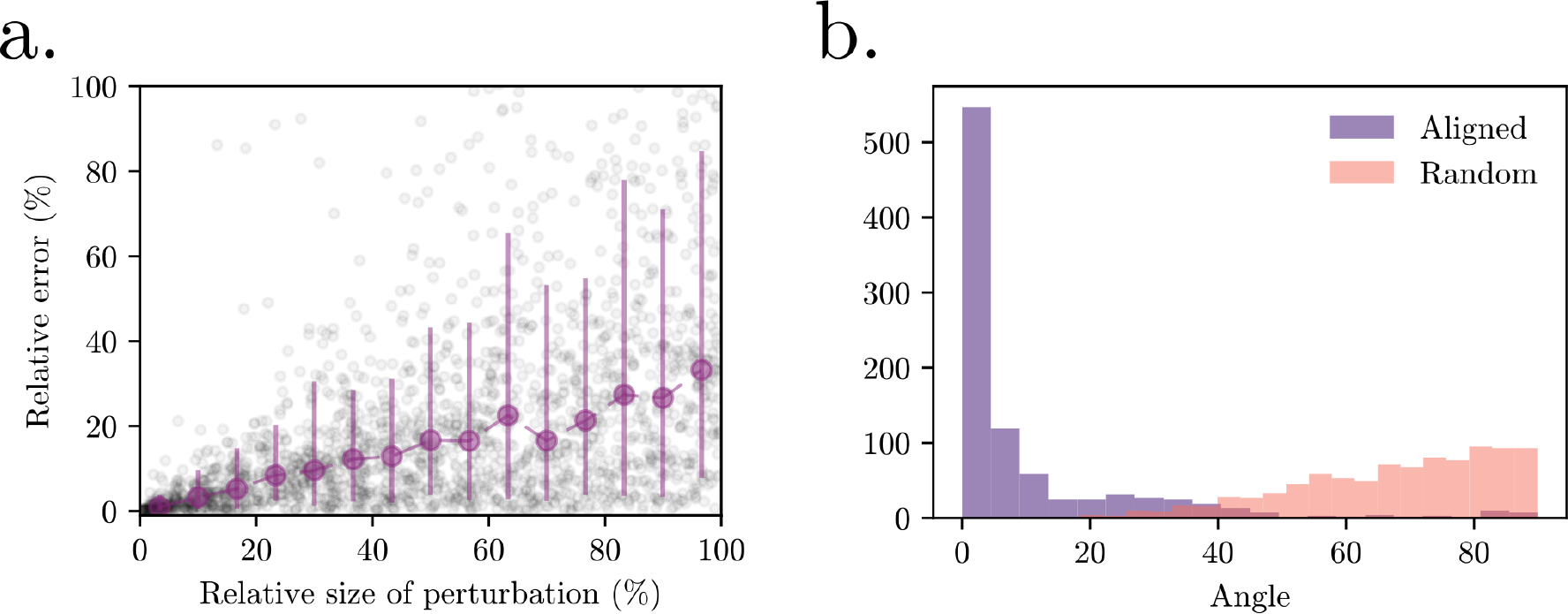
The general framework predicts the effects of mutatations and alignment with environmental perturbations. Panel a. shows the prediction error for the effect of a mutation using the sensitivity vector. The *x*-axis has the perturbation relative perturbation size 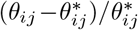. The relative error was calculated as the difference between predicted change using sensitivity vector and the simulated change, divided by the simulated change. 100 random networks were used as reference and 20 mutants were generated for each reference network. Panel b. shows the alignment between genetic and environmental perturbations in *θ*_1,*j*_ and *u*_1_ for random *j* in 1, 2…, 5. The angle between the resulting changes was measured in degrees and plotted as a histogram in purple. Data includes 100 reference networks, each with 20 environmental and 20 genetic perturbations introduced. Orange histogram shows the angle between random vectors in 5-dimensional morphospace.

We now turn to the question of alignment between different sources of perturbation. From the results in *Appendix B*, we know that the sensitivity vectors 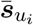 and 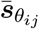 are aligned for any given *i*, and for all *j*. This means that perturbing the environmental parameter *u*_*i*_ or perturbing any of the elements in the *i*-th row of Θ will result in changes in the phenotypes in the same direction. To test this, we generated 100 random reference networks. For each of these reference networks, we introduced genetic and environmental perturbations, in the first row of the reference Θ^*^ and in *u*_1_, respectively. We then simulated the networks until the steady state was reached, and measured the angles between the phenotypic effects of different origins. Figure 3.b. shows that the angles between the vectors resulting from these perturbations are significantly smaller than the angles between random vectors. This confirms that the phenotypic effects of these genetic and environmental perturbations are aligned.

### Plasticity and evolvability

The alignment between the phenotypic effects of genetic and environmental perturbations provides a link between plasticity and evolvability. Indeed, the plastic response of organisms to environmental change can be used to infer what variation can arise through heritable genetic changes, and thus what variation can selection act on.

To study this, we use populations of individuals represented by gene regulatory networks of 5 genes, where only genes 1 and 2 receive environmental input (i.e., *u*_3_ = *u*_4_ = *u*_5_ = 0). We have two sets of 15 populations each that we call *up-down* and *left-right* sets, which differ in how organisms respond plastically to environmental perturbation. Figure 4.a shows one example population from the *up-down* set in black, and one example population from the *left-right* population in orange. The dots represent the steady-states of the phenotypes for the individuals when no environmental input is introduced. The arrows have their origin in these unperturbed reference states, and point in the direction of change when environmental perturbations are introduced. The population plotted in black show large changes in *x*_4_ (i.e., the expression level of gene 4) in response to environmental perturbations, but little change in *x*_3_. The opposite applies to the orange population, which mostly varies in *x*_3_ when environmental perturbations are introduced. Details of how the population sets were generated are given in *Materials and Methods*.

**Figure 4:**
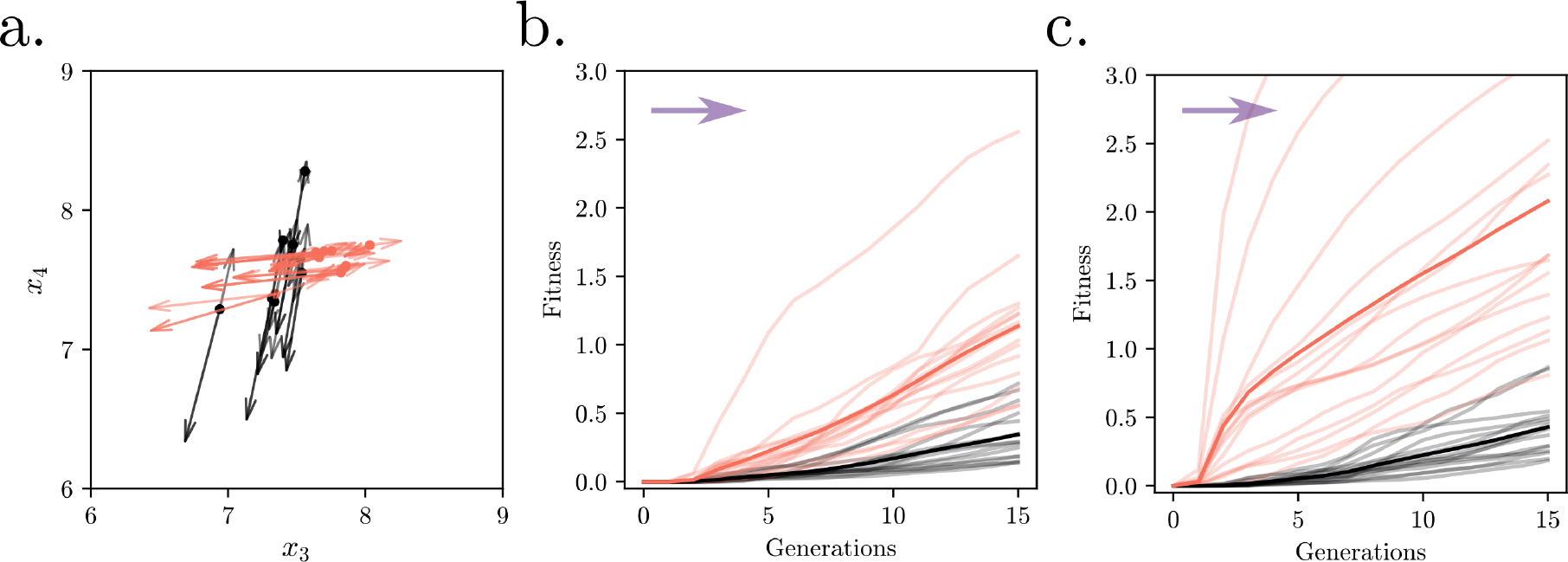
Plasticity predicts evolvability and this can be used for evolutionary control. Panel a. shows an example population of the *left-right* and *up-down* sets, in orange and black respectively. Dots represent unperturbed individuals (no environmental inputs), and the arrows represent the direction of the change in steady states when an environmental perturbation is introduced in one of the first two states. Panel b. shows evolution for *left-right* and *up-down* populations, towards an optimum located in (*x*_3_, *x*_4_) = (12.5, 7.5) represented with the purple arrow. The *left-right* populations, in orange, out-compete the *updown*. Transparent lines are the average among the 25 evolutionary lines initiated from a single individual from each of the 15 populations in each set. Panel c. shows that an additional mutational input directly on the first two rows of Θ significantly accelerates evolution towards the optimum for the *left-right* populations.

Because we know from the analytical results above that the response to environmental perturbations is aligned with the response to genetic perturbations, we predict that the *left-right* populations should evolve faster, compared to the *up-down* populations, if selected in the direction of increase in *x*_3_ and no change in *x*_4_. We test this by making the populations evolve “to the right”, towards an optimum in (*x*_3_, *x*_4_) = (12.5, 7.5). To avoid confounding effects of standing genetic variation, we sampled 25 random individuals from each of the 30 populations (15 *left-right* and 15 *up-down*), and created independent evolutionary lines from 1000 clones of those randomly sampled individuals (i.e., total of 2 × 15 × 25 = 750 independent evolutionary simulations starting from 1000 clones each).

Panel 4.b confirms that the individuals from the *left-right* populations are consistently faster at evolving “towards the right”, to an optimum in (*x*_3_, *x*_4_) = (12.5, 7.5), compared to the individuals from the *up-down* populations. The 15 transparent orange lines correspond to the average of the 25 simulations from each of the 15 *left-right* populations. Similarly, the transparent black lines represent the averages from the 15 *up-down* populations. Total averages are given with fully opaque colors.

For this particular system, we can further use the sensitivity vectors discovered in the previous section to accelerate evolution in a desired direction. From the analytical results, and as confirmed with the simulations (Figure 3.b.), we know that the *i*-th environmental input will be aligned with mutations in the *i*-th row of Θ. Furthermore, we know that the plastic response shown in Figure 4.a. is generated by perturbation in *u*_1_ and *u*_2_. Therefore, we know that evolution in the desired direction can be accelerated by increasing the mutation rate of the first two rows of Θ.

Figure 4.c shows a scenario in which additional mutations are introduced in each generation, but only on the first two rows of Θ. For individuals in set *left-right*, many of these mutations will be beneficial since they will be aligned with the plastic response, which itself points towards the optimum at (*x*_3_, *x*_4_) = (12.5, 7.5), as shown in Figure 4.a. This results in a marked acceleration of evolution towards the optimum (compare orange lines is panels b and c of Figure 4). Populations from the set *up-down*, however, cannot benefit from this additional mutational input since we know from Figure 3.c. that mutations in the first two rows of Θ result in phenotypic changes that do not point towards the optimum for the *up-down* populations. Supplementary Figure 2 shows that, analogously, the *up-down* populations out-compete the *left-right* populations if selection is “upwards”, towards an optimum in (*x*_3_, *x*_4_) = (7.5, 12.5), and that evolution is accelerated in this direction if we increase the mutational input in the first two rows of Θ for the *up-down*, but not the *left-right* population.

## Discussion

In this work, we demonstrate that representing development as a dynamical system provides a theoretical framework to study how generative processes create phenotypic variation, and thus constrain or facilitate adaptive change. This general representation of development captures two general features of generative processes that are lost in static representations that only focus on the outcome of development, but not on how that outcome is constructed (e.g., static maps from genotypes and environments to adult phenotypes). The first property is historicity, which means that the phenotype at any given time is both the effect of earlier, and the cause of later, developmental change. The second property is that all perturbations are ultimately funneled by the same developmental process. These two generic features of development are reflected in the elements of the sensitivity vectors, which determine how the phenotype is expected to change as a result of a perturbation during development.

A benefit of this representation of development as a dynamical system is that it establishes a formal connection between plasticity and evolvability, understood as the capacity to generate phenotypic variation in response to perturbations of environmental or genetic origin, respectively. The existence of theoretical conditions for when genetic and environmental perturbations result in concordant phenotypic effects indicates that both phenomena ought to be more broadly conceptualized as variational properties that reflect the internal structure of the developmental process (Wagner and Altenberg 1996, Salazar-Ciudad 2006). This conceptualization has important implications for evolutionary prediction and control, since it suggests that information of plasticity can reveal salient aspects of evolvability, and vice versa. While this link between plasticity and evolvability has been demonstrated before in specific models (Ancel and Fontana 2000, Espinosa-Soto et al. 2011, Draghi and Whitlock 2012, van Gestel and Weissing 2016, Brun-Usan et al. 2021), the framework presented here extends this understanding in multiple ways.

First, the general framework based on sensitivity functions allows defining explicit theoretical conditions for when plasticity and evolvability should be aligned. These conditions are general and apply to any system of the general form of equation (1), since they are derived from generic features of development represented as a process. Due to their generality, these conditions open the possibility to scale these results for application in evolutionary prediction and control in diverse systems. Importantly, these conditions apply to any point during development and are not constrained to be applied to the adult. This can be important if, for example, selection occurs during development. A limitation for applying this framework to phenotypic variation in nature is that the developmental function ***f*** needs to be known to calculate matrix *A*(*t*) and vector *b*(*t*). Note however that even if we cannot obtain explicit analytical values for these elements, the general conclusions of the framework still apply. Furthermore, there is potential to estimate the sensitivity functions directly from data when the developmental function is not known (e.g., by analyzing the phenotypic consequences of experimental perturbations, Milocco and Uller 2023).

Second, the framework makes it possible to go beyond the qualitative expectation that genetic and environmental perturbations are interchangeable in development (e.g., Cheverud 1982, 1988; West-Eberhard 2003), by making quantitative predictions of the phenotypic effect of perturbation, and of the relationship between different perturbations. As shown in this paper, this information can be exploited to predict responses to selection without an estimate of heritable (co)variance in phenotypes (e.g., as summarized in the G matrix). This is possible because knowledge about plasticity captures properties of developmental systems that carries information about how those systems can accumulate heritable phenotypic variation. While some empirical data (e.g., Noble et al. 2019) could be interpreted in this manner, there appears to be no direct test of this prediction. Note, however, that the framework introduced here is only locally valid, meaning that it is predictive of the effects of perturbations of small size. Therefore, predictions of evolvability based on plasticity may only be valid for a limited number of generations after which the sensitivity functions would have to be re-identified since the internal structure of development may have changed.

Finally, the framework reveals how to exploit this alignment for evolutionary control, by accelerating evolution in certain directions through forced mutations predicted to result in adaptive phenotypic changes. More generally, if the sensitivity vectors of multiple perturbations are identified, this means that it is possible to design combinations of perturbations to drive the developmental process in a desired direction. Similar to the other points above, this insight suggests opportunities for empirical investigation of evolvability, which also may have implications in applied fields of biology such as biotechnology.

While so far we have emphasized the alignment between the phenotypic effects of genetic and environmental perturbations, different genetic perturbations can also be aligned with each other (Pitchers et al. 2019). In the simulations, this redundancy is evidenced by the fact that mutating any element of the *i*-th row of Θ in the gene regulatory network or mutating any of the parameters *k* or *F* in the reaction-diffusion model, generates concordant phenotypic change. In this way, a population will evolve in the same direction of trait space by accumulating mutations in any of those equivalent elements. This redundancy can explain why genetic changes underlying parallel evolution often fail to be replicated (e.g., Pelletier et al. 2023), since multiple changes at the genetic level can explain the same phenotypic adaptations.

Redundancy ultimately reflects the fact that it is not the identity of any specific gene that matters for the generation of phenotypic variation, but rather the role it plays in the dynamics of the developmental process. This can explain the observation that, despite the multidimensional nature of phenotypic data, it is very often the case that there are only a few effective dimensions of variation (Beldade et al. 2002, Houle et al. 2016, Alba et al. 2021, Rohner and Berger 2023). Indeed, if many perturbations result in concordant phenotypic changes, then phenotypic variation will be restricted to a manifold of lower dimension than the total number phenotypic variables.

Following this, we should expect that parallel evolution will be explained by a repeatable genetic change only in cases where the effect of perturbing that gene is unaligned with others (i.e., the associated vector 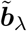 in equation (5) is unique). This represents a scenario where the perturbed gene plays a distinctive role in developmental dynamics. One possible example of this is the gain and loss of red and yellow carotenoid coloration in diverse vertebrates (e.g., birds, mammals, lizards), which is commonly associated with perturbations in the expression of gene *BCO2* that encodes a carotenoid degradation enzyme (Våge and Boman 2010, Andrade et al. 2019). This evidence implies that *BCO2* plays a distinctive role in the generation of color, so that perturbations in its functioning have distinctive phenotypic effects. Repeatable genetic changes underlying parallel evolution can thus be used to make inferences about developmental dynamics, guiding future research.

## Conclusion

An understanding of evolution is incomplete without a theory of how phenotypic variation is generated in each generation. The representation of development as a process provides the conceptual basis to predict when perturbations of different origins result in similar phenotypic changes. Our results indicate that a promising avenue for future research on the generation of variation will not focus on the specific identity of elements such as genes, but rather focus on how those elements participate in a dynamical process that integrates different sources of information to produce phe-notypes.

## Materials and Methods

### Reaction-diffusion simulations

The tissue is composed of a grid of 50 × 50 cells, each having a given amount of the two morphogens, thus resulting in a total of 5000 states. Difussion occurs only between neighboring cells and is represented with a discretized version of the Laplacian as commonly done (e.g., Sick et al. 2006). Periodic boundary conditions are assumed, so the tissue can be thought of as being mapped to a torus. The first 2500 states are the concentration of morphogen 1 in the 2500 cells, while the last 2500 states correspond to the concentration of morphogen 2. This means that for *i* = 1, 2, …, 2500, *x*_*i*_ corresponds to concentration of morphogen 1 of the cell located in position (*q* + 1, *r*) of the grid where *q* and *r* are the quotient and remainder, respectively, of the division *i* ÷ 50, where ÷ represents integer divison. Similarly, *x*_*i*_ for *i* = 2501, 2502, …, 5000 corresponds to the concentration of morphogen 2 of the cell located in position (*q* + 1, *r*) of the grid where *q* and *r* are the quotient and remainder, respectively, of the division (*i* − 2500) ÷ 50. To obtain the sensitivity vectors, we need the Jacobian and the weights. These are obtained by differentiating the discretized equation. The Jacobian results in a sparse matrix since only neighboring cells interact (through diffusion). The Jacobian and the weights are given in the accompanying scripts.

#### Populations of gene-regulatory networks

To generate the *up* − *down* and *left* − *right* populations, composed of individuals that have plastic responses in different directions, we evolve populations under fluctuating selection tracking an optimum with a correlated environmental input (see Draghi and Whitlock 2012). Specifically, in each generation, the position of the optimum is correlated with the environmental inputs, so the networks with higher fitness are those that allow individuals to tract the optimum in each generation using the environmental input. To simplify the figures, out of the 5 genes, only genes 1 and 2 receive environmental input and selection acts only on genes 3 and 4. We evolve the populations for 500 generations, starting with 1000 clones that were randomly generated. All optimums fluctuated around a value of (*x*_3_, *x*_4_) = (7.5, 7.5). A set of 15 populations were evolved to track an optimum that fluctuated for values of *x*_3_ around 7.5, but keeping *x*_4_ fixed at 7.5. We refer to this first set of populations as *left-right*. Another set of 15 populations, which we refer to as *up-down*, evolved tracking an optimum where *x*_4_ fluctuated and *x*_3_ was fixed at 7.5.

#### Angle between sensitivity vectors

The angle *θ*(*t*) between the directions of two sensitivity vectors 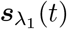 and 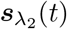 is defined as the minimum between the angle formed by 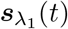 and 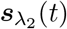, and the angle formed by 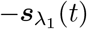 and 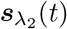. By taking the minimum we make sure that *θ*(*t*) depends on the direction of the vectors and not the sign. Complete alignment between 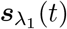 and 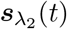 is then given by *θ*(*t*) = 0°, in which case the sensitivity vectors have the exact same direction (but possibly different signs). This will be the case if the weights are proportional (i.e., 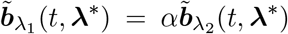 with *α* constant). More generally, however, we consider that there is evidence for alignment if the angle *θ*(*t*) is significantly smaller, at a given confidence level, than the distribution of angles between independent random vectors in ℝ^*n*^, with *n* being the number of states. We note that as *n* becomes larger, the distribution of angles between random vectors in ℝ^*n*^ concentrates around 90°, so random vectors are generally ‘more’ orthogonal in higher dimensional spaces. In this way, *θ*(*t*) *<* 90° is higher dimensions is a signature of alignment.

## Appendixes

### Appendix A

Here, we obtain an expression for the sensitivity vectos. We begin with the general representation of development as a dynamical system, as given by equation (1). Under the assumption that the developmental function ***f*** is smooth, there exists a unique solution ***x***(*t, λ*) for a *λ* close to the reference value *λ*^*^, which can be obtained as (see Khalil 2002)

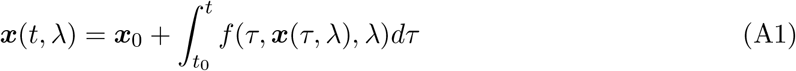

We are interested in calculating the sensitivy vector, as defined in equation (2). For this, we take the partial derivative with respect to *λ* on both sides of equation (A1), which gives

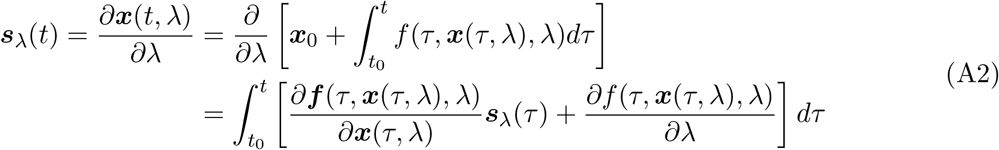

where we use that that the derivative of the integral is equal to the integral of the derivative, and that the derivative of the initial condition is zero because it does not depend on *λ*. We further use the chain rule to obtain the derivative of ***f*** with respect to *λ*.

To obtain an expression of how the sensitivity vector changes in time, we take the partial derivative of the equation (A2) with respect to time, yielding equation (4).

### Appendix B

Here, we derive the equations for the Jacobian, weights and sensitivity vectors for the developmental parameters of the gene regulatory network. We begin by rewriting equation (7) in matrix form. For this, we write ***h*** = Θ***x*** + ***u***, and define ***κ*** = (*K*_1_, *K*_2_, …, *K*_*n*_) and the *n* × *n* diagonal matrix *M* = *diag*(*μ*_1_, *μ*_2_, …, *μ*_*n*_). This yields

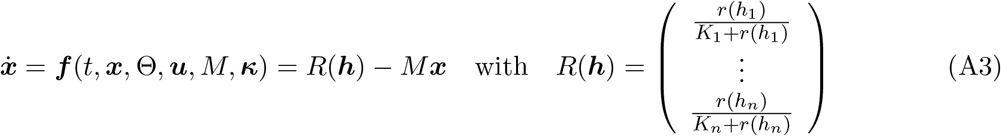

which follows the form of equation (1), with the developmental parameter vector given by ***λ*** = (Θ, ***u***, *M*, ***κ***) and reference developmental parameter values given in the vector ***λ***^*^ = (Θ^*^, ***u***^*^, *M* ^*^, ***κ***^*^).

The study of alignment in this example is performed at the steady state, with 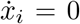 for all *i*. We therefore assume that the reference values for the developmental parameters, given by ***λ***^*^, result in a stable system able to reach a steady state. Further, we assume that 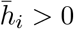, with the bar indicating the steady state. This allows to replace 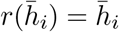 Note that if 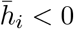, then equation (7) reduces to 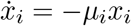 and it can be easily checked that this results in 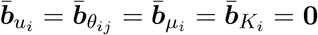.

In other words, the steady state is robust to perturbations in the developmental parameters if 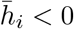.

We obtain the Jacobian,

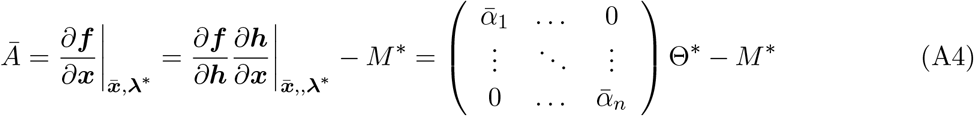

where the bars indicate variables in the steady state, asterisks indicate reference value, and 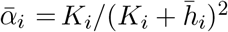. We now calculate the weights 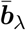as

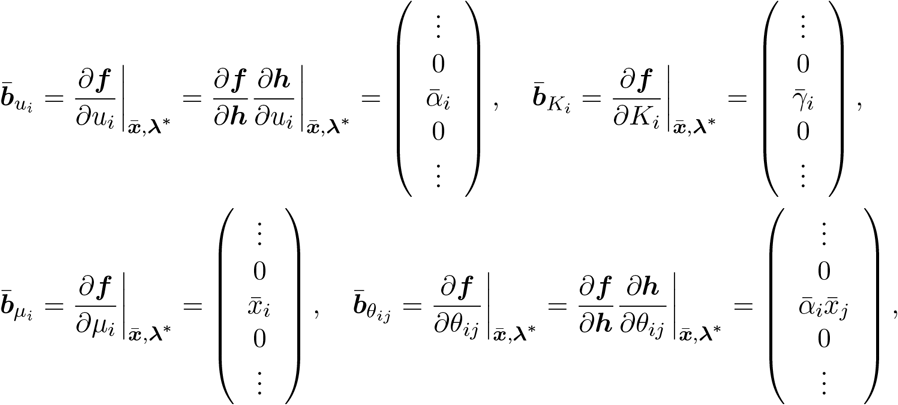

with 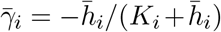. Because the Jacobian is invertible, we can obtained the sensitivity vectors with the simplified expression given in equation (5), as

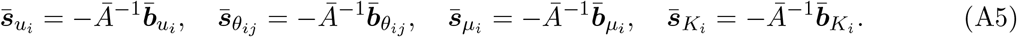

Thus, the sensitivity functions for *u*_*i*_, *θ*_*ij*_, *μ*_*i*_, and *K*_*i*_ are always aligned for a given *i* and all *j*.

In particular, they point in the direction of the *i*-th column of *Ā*^−1^.

## Acknowledgements

The authors thank Ruben H. Milocco for discussion. The authors thank the John Templeton Foundation (62220) for financial support. The opinions expressed in this paper are those of the authors and not those of the John Templeton Foundation.

## Data availability

The *Python* scripts for the reaction-diffusion and gene regulatory network simulations and analyses will be uploaded to https://github.com/lisandromilocco.

## Supplementary Figures

**Supplementary Figure 1.**
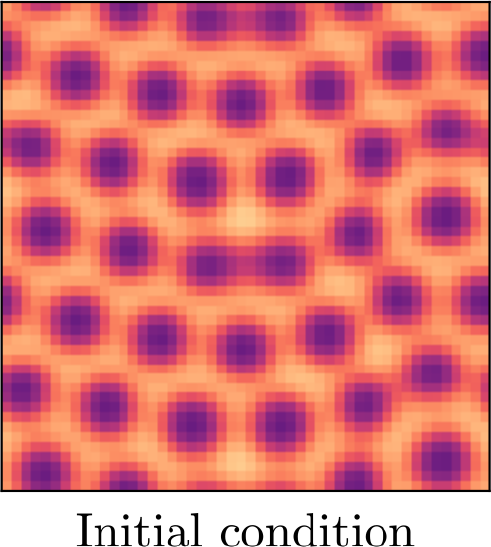
Initial conditions for the reaction-diffusion simulations.

**Supplementary Figure 2.**
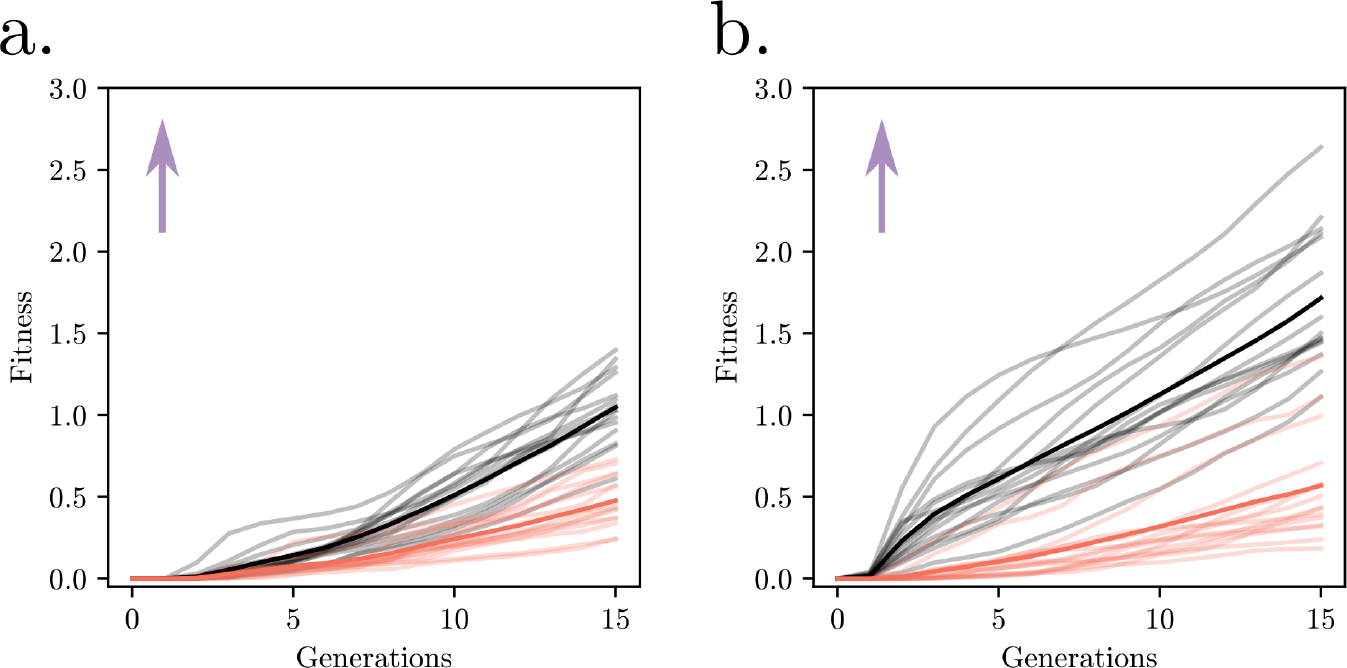
This figure is analogous to Main Figure 4 but for selection “upwards”, towards an optimum in (*x*_3_, *x*_4_) = (7.5, 12.5) as represented by the purple arrow. Panel a. shows evolution for *left-right* and *up-down* populations, in orange and black respectively. The *up-down* populations out-competes the *left-right*, as expected from their plastic responses shown in Main Figure 4.a. Transparent lines are the average among the 25 evolutionary lines initiated from a single individual from each of the 15 populations in each set. Panel b. shows that an additional mutational input directly on the first two rows of Θ significantly accelerates evolution towards the optimum for the *up-down* populations.

